# A novel pipeline for the rapid expansion of ecological trait databases using LLMs

**DOI:** 10.64898/2026.03.10.710865

**Authors:** Robert J. Ramos, Michelle E. Afkhami, Carlos A. Aguilar-Trigueros, Kristin M. Barbour, Priscila Chaverri, Sarah A. Cuprewich, Cameron P. Egan, Kira M. T. Lynn, Kabir G. Peay, Veera Norros, Adriana L. Romero-Olivares, Lauren Ward, Bala Chaudhary

## Abstract

This paper presents a novel workflow leveraging Large Language Models (LLMs) to rapidly extract trait data from fungal species descriptions, addressing a significant bottleneck in ecological research. We developed and evaluated an LLM pipeline to extract morphological trait data from arbuscular mycorrhizal fungi, comparing performance against a manually curated dataset (TraitAM). Results demonstrate the potential of LLMs for automated trait data acquisition, though accuracy varies by trait and model, with systematic biases observed. This framework offers a blueprint for building trait databases across diverse taxa and domains, significantly accelerating ecological research and conservation efforts.

## Introduction

The utility and importance of using trait data (i.e. measurable characteristics of biological species that affect its fitness or ecological performance) continues to attract attention across ecological fields^1,2^. These trait data are paramount for the development of predictive models to understand the underlying mechanisms of the response of biodiversity to global change across all domains of life, from plants to microbes^3^, vertebrate mediated environmental services^4^, predicting fungal responses to environmental stress^5^, and determining colonization abilities in microbes^6,7^. The major bottleneck of predicting functional frameworks for biodiversity is the availability of comprehensive trait data for taxa or populations of interest^8^. While functional trait databases with searchable fields do exist ^9–12^, they are fragmented both in terms of trait and species coverage. Paradoxically, there exists an abundance of usable data that could be fed to these databases, but it is concealed in complex text resources (e.g. taxonomic descriptions). With several thousands of scientific papers and vast taxonomic literature where these data are found, manually extracting trait data is a time-consuming and error-prone process. To solve this limitation, we propose a workflow that utilizes state-of-the-art Large-Language Models (LLMs) to automate the process of parsing species descriptions and generating structured trait datasets.

Computational analytics hold the potential to transform humanity’s ability to understand enigmatic biodiversity, improve conservation and management of natural systems, and tackle long-standing questions critical to environmental and human health. Research leveraging large, publicly-assessible datasets has already revealed new insights into alternative lifestyle strategies of the bacteria and fungi that maintain soil health^3^, and conservation of biodiversity^13,14^. The volume of environmental data is expanding rapidly through several pathways. First, technological advancements now produce orders of magnitude more data at a fraction of the former cost or effort^15–17^. Second, large-scale digitization initiatives have significantly enhanced the global accessibility of rich historical and newly-acquired data that once were confined to a particular location^18,19^. Despite these advances, a great deal of “available” data is not in a usable form for addressing the important fundamental and applied questions they could inform^20^. This paradox of accessibility without usability captures a defining tension of modern science, which must be resolved to harness Big Data for ecological and societal progress^20,21^. This challenge is exemplified in the proliferation/abundance of biological information in unstructured textual form (e.g. taxonomic descriptions), which represents a significant bottleneck for ecological modeling, conservation efforts, and biodiversity research^22^.

The speed and automation associated with Artificial Intelligence (AI) offers advantages for extracting valuable data from dense textual resources. What is AI can be a difficult concept to pin down. A useful definition is that of a rational agent that “should do whatever action is expected to maximize its performance measure, on the basis of the evidence provided by the percept sequence and whatever builtin knowledge the agent has”^23^. AI techniques to extract structured data from text is not new. Natural language processing (NLP) techniques and pipelines have existed for specialized data extraction tasks since the first demonstration of machine based translation in 1954^24^. Early models were rules based, attempting to capture the meaning and semantics of language in predefined rules. In the 1990s this approach gave way to statistical models that used machine learning techniques to train on large corpura^25^. Since 2015 traditional statistically informed models have been outperformed by neural networks which can utilize a multitude of algorithms including transformer models which form the foundation of modern LLMs^26,27^.

Large Language Models (LLMs) have the potential to automate the tedious process of manually sifting through text from scientific papers to extract needed information about the ecology of thousands of species gathered by the scientific community. For instance, organismal attributes and responses of organisms (e.g. physiological, behavioral, phenological, etc.) to experimental treatments^28,28,29^, and morphological, physiological or behavioral traits. LLMs are machine learning models constructed from a transformer neural network architecture and trained on massive datasets of human-generated text^26^. These models, which are trained adjusting millions or even billions of parameters to minimize prediction errors, learn to predict the probability of the next word given the preceding words. This approach effectively builds a complex statistical representation of language structure, semantics, and ontologies^30^, and subsequently, these models can be used to generate text, translate languages, and answer questions. Consequently, LLM automation has the potential to dramatically reduce the time and effort required to answer a diversity of research questions that require data compilation that was previously prohibitively time-consuming.

These recent technological advances represent a major leap in our ability to build trait databases, particularly for groups where trait information is scarce or absent. Yet, because these approaches constitute a fundamental departure from previous technologies, these models necessitate not only new workflows but also a careful assessment of their strengths and limitations. We demonstrate the power of LLMs by extracting trait data for arbuscular mycorrhizal fungi, a group of symbiotic fungi associated with over 70% of plant species and of major ecological and economic importance worldwide. This group provides an ideal test case for two reasons. First, despite their importance, trait data for arbuscular mycorrhizal fungi remain scarce, limiting our ability to build predictive ecological models. Second, a recently published database of spore traits compiled through manual extraction from taxonomic descriptions (TraitAM) offers a unique opportunity to benchmark automated LLM approaches against existing data^12^. Specifically, we developed an LLM pipeline designed to identify and extract quantitative and qualitative trait data (spore size, spore wall thickness, and ornamentation height) from species descriptions found in publicly available databases and scientific publications. We evaluated the effectiveness of 2 LLM models across three overall approaches (Figure 1). The traitAM dataset allowed us to judge reproducibility, fidelity to expert derived values, and systematic bias in our approach. This also provided valuable insights on where LLM based approaches still need to improve. The development of this robust LLM pipeline represents a significant advancement in computational methods for biological data curation. Overall we find that LLM trait extraction can achieve similar performance to expert extracted values, but this was highly variable between traits. Expert supervision is recommended to ensure LLMs are performing as expected. In addition, by automating data extraction we ensure a more efficient and reproducible scientific process. Altogether, this workflow emphasizes repeatability of outputs, reproducibility of the entire pipeline, and evaluation of efficacy with a reference dataset. It offers a blueprint to apply LLMs to build trait databases across new traits (growth form, fruiting body height, habitat preference, among many others) and all domains of life.

**Figure 1.**
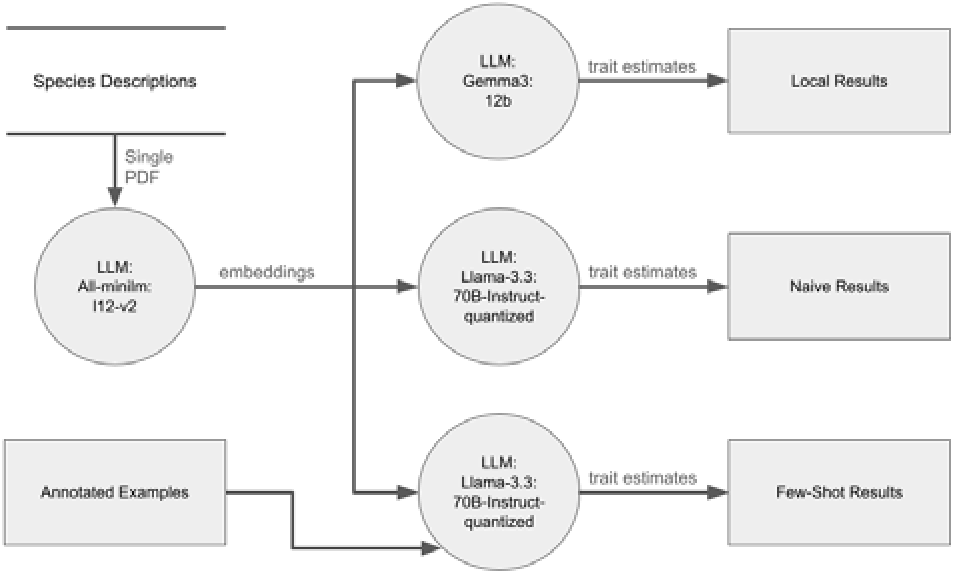
Data flow diagram of LLM generation of trait data. This diagram gives an overview of the pipeline used to generate the results for our three approaches (Local, Naive, Few-Shot). The open box is our data store of pdfs. Closed boxes are input or outputs, and circles represent processes (i.e. our LLM models).

## Methods

Preparation of this text was assisted with the use of LLMs, hosted locally with Ollama platform^31^. Gemma 3^32^ was used to enhance readability and language, and in formulating and restructuring content. The authors have reviewed and edited all generated text and take full responsibility for its content. To develop and evaluate our workflow for extracting fungal trait data, we followed a systematic approach that involved data collection, model implementation, and iterative refinement. Below, we describe each step in detail.

### Data Collection

The first step in our workflow involved gathering a comprehensive dataset of fungal species descriptions. We sourced these descriptions from the TraitAM database, a publicly available repository of spore trait data for arbuscular mycorrhizal fungi^12^. We used the descriptions of TraitAM because it allows us to directly compare the trait values extracted with our LLM pipeline to the ones extracted manually by the experts for TraitAM. The database provided access to a wide range of scientific publications and species descriptions in PDF format as well as trait data manually extracted from those documents by experts. These documents served as the primary input for our pipeline. Each PDF contained detailed information about a fungal species, including both quantitative traits (e.g., spore size, spore wall thickness) and qualitative traits (e.g., ornamentation type). Data on spore size, spore wall thickness, and ornamentation height had been extracted by experts for TraitAM and were therefore available as expert provided comparison values.

### Data Ingestion and Preprocessing

Once the PDFs were collected, we ingested them into the pipeline using a Retrieval-Augmented Generation (RAG) framework. Each document was processed individually, limiting the number of species each LLM would have to process at a time. An important limitation of the current pipeline is that it does not have a dedicated mechanism to handle documents that describe multiple species, and will describe only a single species from those documents. The RAG framework allowed us to combine the strengths of information retrieval and generative language modeling, enabling the pipeline to focus on relevant sections of the text while minimizing noise from unrelated content. This step was completed using the All-minim model, which was specifically trained for embedding tasks^33^. Preprocessing steps included text extraction and formatting to ensure compatibility with the downstream language model (Figure 1).

### Model Implementation

We crafted prompts for the models to extract and assign for each species values for the following spore traits: length, width, minimum wall thickness, maximum wall thickness, minimum ornamentation height, and maximum ornamentation height. As mentioned above, we target these traits because they could be directly compared to values manually extracted by experts for the TraitAM database. Our initial implementation utilized a local instance of the Gemma 3: 12 billion parameter model, which was run on the Ollama platform^31^. Temperature was set to 0.7, context size was set to 48000, and streaming was disabled. The limit on retries was increased to 25 and the model was run 10 times with a series of static seeds (42, 1337, 2020, 7, 469, 6150, 8611, 8775, 7952, 2885, 2885). Gemma 3 is a state-of-the-art large language model designed for natural language understanding and generation^32^. The local deployment allowed us to maintain control over the data and ensure compliance with data privacy requirements. As the project progressed, we transitioned to using the Llama 3.3: 70 billion parameter instruct model, hosted on the CyVerse Verde platform^34,35^. Again, temperature was set to 0.7, and the limit on retries was increased to 25. The model was run 10 times with a series of static seeds (42, 1337, 2020, 7, 469, 6150, 8611, 8775, 7952, 2885, 2885). This model represents a larger and more advanced language model than could be run on local hardware, with 70 billion parameters fine-tuned for instruction following tasks. The CyVerse Verde platform provided the computational resources necessary to handle the increased complexity and scale of the model. In our results we present comparisons between outputs of the two LLM models. This transition significantly improved the accuracy and efficiency of the data extraction process, particularly for more nuanced or context-dependent traits.

### Few-Shot Training and Refinement

To further refine the model’s performance as measured by percent deviation from expert derived values, we employed a few-shot training approach. This involved providing the same Llamma 3.3-70b-Instruct model with a small number (3) of single species annotated examples from different genera (*Acaulospora* and *Gigaspora*) to guide its understanding of the task (Supplement 1). These examples were carefully curated to represent the diversity of fungal traits and the variability in how they are described in the literature. The few-shot training process allowed the model to generalize from the examples and improve its ability to extract relevant data from unannotated texts. The performance of this model was compared to the “naive” Gemma3 and Llama3 model results which were not given annotated examples.

### Measuring Performance

Model performance was measured as the percent difference between LLM outputs and expert derived values for each species according to equation 1,

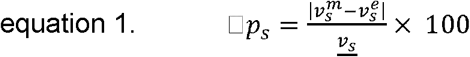

Where for each species *s*, 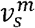 is the model provided value, 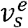 is the expert provided value, and *<u>v</u>*_*s*_ is the mean between them. Expert provided values for spore width were the mean of dim2.min, dim2.min2 dim2.max, dim2.max2 from TraitAM. Expert values for length were the mean of dim1.min, dim1.min1 dim1.max, dim1.max2. Expert values for minimum and maximum wall difference were taken directly from the traitAM min_wall_thickness and max_wall_thickness respectively. Similarly, expert values for minimum and maximum ornamentation height were taken from the traitAM orn_height_min and orn_height_max respective values.

Statistical analyses were conducted using the StatsModels^36^ and MarginalEffects^37^ python packages. ANOVA analysis was conducted on LLM values predicted by approach (Local, Naive, or Few-Shot) and replicate (statsmodels.formula.api.ols and statsmodels.stats.anova_lm). Ordinary least squares (statsmodels.formula.api.ols) were conducted for each trait on expert values predicted by LLM Values by approach. Hypothesis testing for H_0_ = 1 was conducted using the MarginalEffects package (marginaleffects.hypothesis). Finally, a Generalized linear model (statsmodels.formula.api.glm) analysis was conducted on percent differences predicted by replicate run and approach interacting with traits. The family was set to Tweedie (statsmodels.families.Tweedie) using a variance power of 1.1.

## Results

We evaluated the performance of three LLM-based approaches for extracting morphological trait data from fungal species descriptions: a local naive (zero-shot) Gemma3:12b model, a naive Llama-3.3-70B implementation, and a few-shot trained Llama-3.3-70B implementation.

Each approach was tested with 10 replicate runs using different random seeds to assess extraction consistency and accuracy across six fungal trait categories: spore length, spore width, minimum wall thickness, maximum wall thickness, minimum ornamentation height, and maximum ornamentation height.

### Extraction Accuracy Across Approaches and Traits

Percent differences between LLM-extracted and expert-derived trait values varied substantially across both extraction approaches and trait categories (Table 1). The local Gemma3:12b model consistently produced the largest deviations from expert measurements across all trait categories, with a mean percent difference of 65.08 (57.65) compared to 49.75 (62.44) and

**Table 1.** Anova Results.

| Trait | Model Term | Sum of Squares | df | F | Pr(>F) |
| --- | --- | --- | --- | --- | --- |
| Length | approach | 237128 | 2 | 9.376 | 0 |
|  | run | 2619 | 1 | 0.207 | 0.649 |
|  | Residual | 87495531 | 6919 |  |  |
| Width | approach | 1841761 | 2 | 89.504 | 0 |
|  | run | 43 | 1 | 0.004 | 0.949 |
|  | Residual | 70858377 | 6887 |  |  |
| Minimum Wall Thickness | approach | 2700 | 2 | 155.895 | 0 |
|  | run | 7 | 1 | 0.785 | 0.376 |
|  | Residual | 50879 | 5875 |  |  |
| Maximum Wall Thickness | approach | 14339 | 2 | 15.034 | 0 |
|  | run | 204 | 1 | 0.428 | 0.513 |
|  | Residual | 2890393 | 6061 |  |  |
| Ornamentation Height Min | approach | 1018 | 2 | 171.36 | 0 |
|  | run | 0 | 1 | 0.053 | 0.819 |
|  | Residual | 8775 | 2954 |  |  |
| Ornamentation Height Max | approach | 769 | 2 | 17.656 | 0 |
|  | run | 16 | 1 | 0.717 | 0.397 |
|  | Residual | 64315 | 2953 |  |  |

51.51 (55.68) for the few-shot model and naive Llama3 models respectively. None of the LLM models or Traits showed statistically significant variation between individual runs (Table 1).

Spore dimensional measurements (length and width) exhibited the lowest percent differences between LLM estimates and expert obtained values across all three approaches, with median differences below 25% for both the naive and few-shot Llama-3.3 implementations (Figure 2). In contrast, ornamentation height measurements showed the greatest variability in extracted trait values between model runs and highest percent differences with expert values, particularly for minimum ornamentation height. Wall thickness measurements showed intermediate performance, with the few-shot approach demonstrating notably reduced variability compared to both the local and naive approaches.

**Figure 2.**
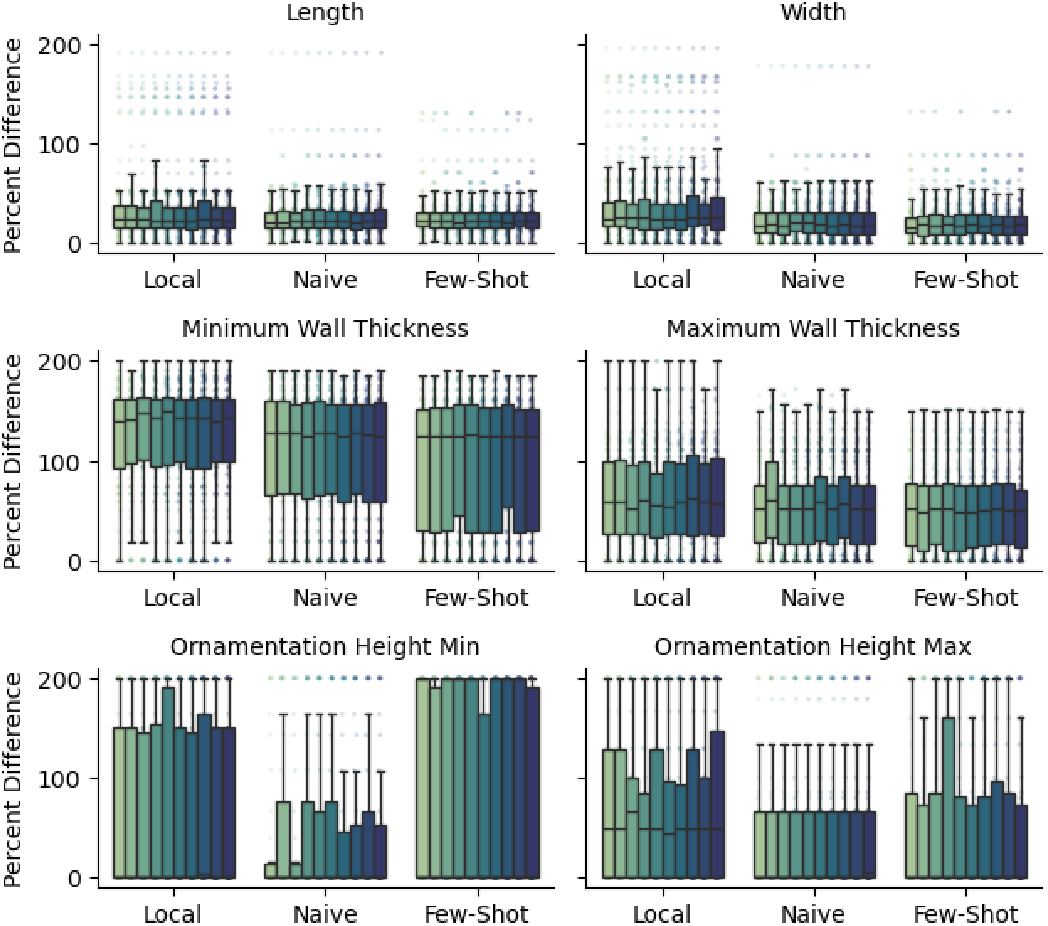
Percent difference by individual replicate run. Percent differences of LLM values to expert extracted values (TraitAM) are plotted by fungal trait. Each bar represents an individual replicate run (1-10) for each extracted trait. Results are grouped by approach, with the local Gemma3 model as the leftmost group, the naive Llama3 results as the center group, and the few-shot trained Lamma3 results as the rightmost group. Percent difference was smallest for spore width and length. The greatest variability was seen with minimum and maximum ornamentation height.

### Systematic Bias in LLM Trait Extraction

Correlation plots between LLM-extracted and expert-derived values revealed systematic biases in trait measurement across approaches (Figure 3). The local Gemma3 model demonstrated consistent underestimation across all trait categories, with regression lines falling substantially below the 1:1 reference line. This suggests the local model struggled to accurately parse numerical trait information from species descriptions. Both Llama-3.3 approaches showed improved fidelity relative to expert values, with regression lines more closely approximating the 1:1 relationship. The only regression lines whose slopes were not statistically different from 1 was the Few-Shot model for minimum ornamentation height (p=0.275) and the naive model for maximum ornamentation height (p=0.704) (Supplement 2).

**Figure 3.**
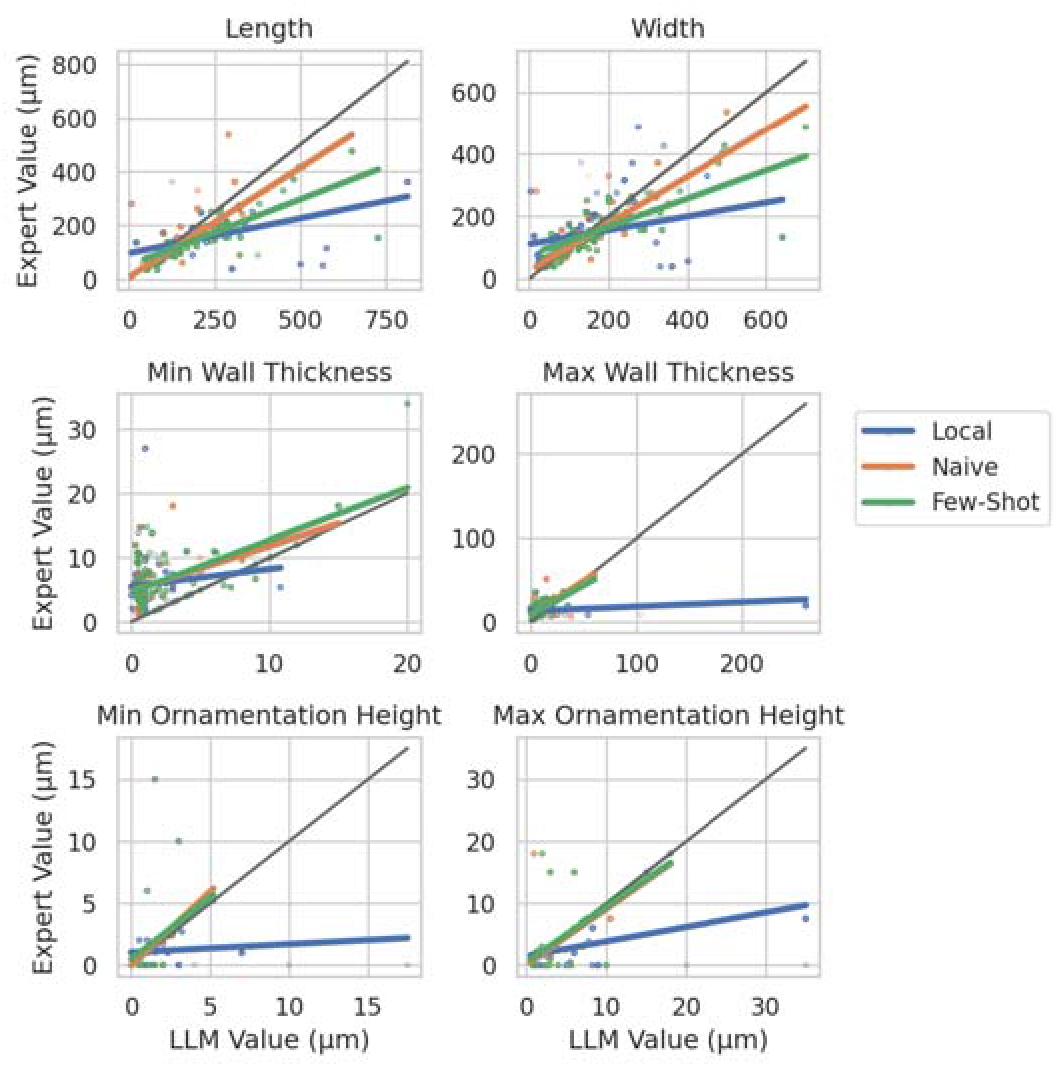
Correlation of LLM and expert derived values. Expert derived values for each trait (in μm) is plotted on the y-axis with the LLM derived values on the x-axis. Plots are separated by fungal traits. Individual species data points are represented as colored dots with simple regression lines also plotted. A black 1 to 1 line is also presented for reference. For all traits the local Gemma3 model has the greatest bias, with the LLM underestimated values.

### Statistical Comparison of Extraction Approaches

A generalized linear model (Tweedie family, variance power = 1.1) quantified differences in extraction performance across approaches while controlling for trait type and replicate run effects. Model-predicted percent differences confirmed significant main effects of both extraction approach and trait category, as well as a significant interaction between these factors (Figure 4).

**Figure 4.**
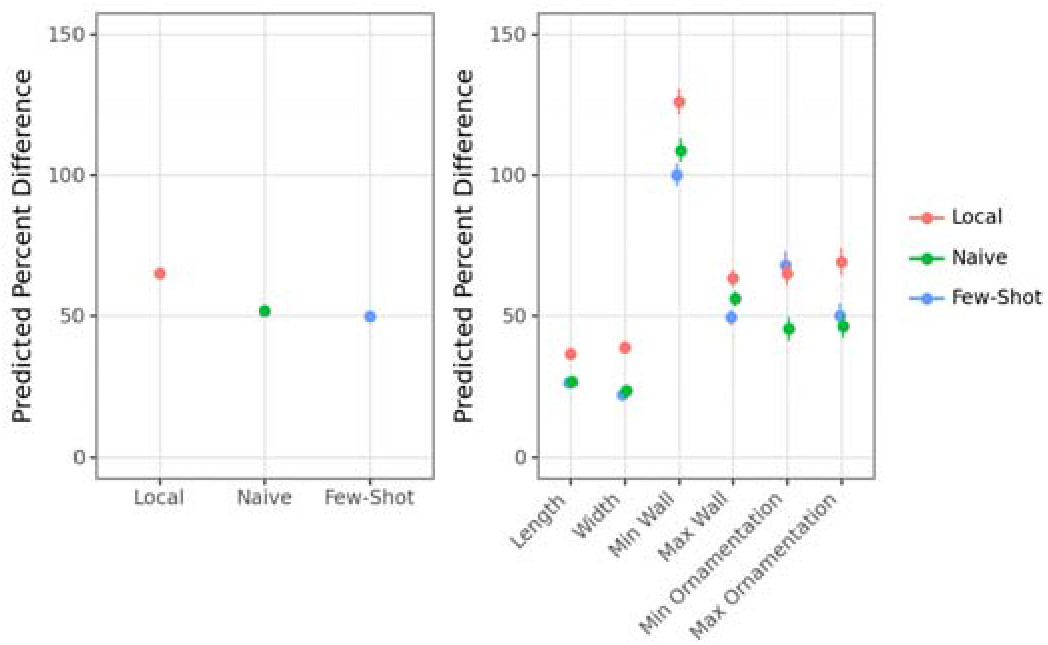
Marginal effects of predicted percent difference. Marginal effects from a generalized linear model (Tweedle family, variance power = 1.1). Marginal effects for both main effects of approach (a) and the interaction of approach with fungal trait (b).

The local approach had statistically significantly greater predicted percent differences compared to both naive (p < 0.001) and few-shot (p < 0.001) approaches, with predicted values approximately 14% higher than the naive approach and 15% lower than the few-shot model (Figure 4a). The few-shot model did not have statistically significantly lower percent difference than the naive model (p =0.217), demonstrating that there was not an improvement in data extraction precision when the model was given annotated examples.

The interaction analysis (Figure 4b) revealed that the benefits of few-shot training varied by trait category. The approach-by-trait interaction showed the greatest improvements for wall thickness measurements, where the few-shot approach reduced predicted percent differences by 9% for minimum wall thickness and 7% for maximum wall thickness (p=0.003 and p=0.001 respectively). For the remaining spore dimensions, percent difference was not significantly reduced in the few-shot model for either length or width (p=0.802, p=0.276). Ornamentation height measurements didn’t see a benefit for the few shot model for maximum ornamentation size (p=0.184), with the naive model having a statistically lower percent difference than the few shot model by 22% (p < 0.001).

## Discussion

We were able to successfully use our novel pipeline to extract arbuscular mycorrhizal trait data from species descriptions. For certain traits we were able to achieve similar performance to TraitAM (<25 percent difference). With other traits, such as minimum and maximum spore wall width, there was much greater deviation from the expert derived values. This highlights the need for expert supervision to ensure models are behaving as expected. However, provided adequate safeguards, this method shows promise in accelerating trait based research.

Large Language Model size (12 billion parameter Gemma 3 vs the 70 billion parameter Llama 3.1 model) corresponded with improved accuracy of trait extraction, and this was true averaged across all traits (Fig 4a). The larger Llama (both Naive and Few-Shot) models improved accuracy for all traits individually except minimum ornamentation size. This result aligns with previous research showing a positive relationship with overall model size and performance ^38^. Importantly model size has been shown to have counter active effects on overall performance, with larger models showing greater “informativeness”, while smaller models are more likely to have increased “faithfulness” ^39^. Here informativeness is a metric of the models ability to transform essential information into an appropriate form^40^; however, faithfulness tracks a model’s propensity for not generating inaccurate or irrelevant facts^40,41^. There is a tradeoff present for model size and this is before considering the computational and energy costs of running a model larger than the task necessitates. Therefore careful consideration should be placed on model size selection for a particular analysis.

LLM accuracy varied between different traits, with more complex traits decreasing the overall effectiveness of all three approaches. Spore length and width were extracted with the greatest overall accuracy (smallest percent difference from expert values) and minimum wall thickness had the lowest. Wall thickness is a trait that often needs to be calculated from descriptions. AMF may have multiple layers to their spore wall (1-4) and that can be an important detail for the species descriptions. This means for many species the LLM had to combine different thickness metrics. LLMs are particularly poor at mathematical operations, and this may account for the higher inaccuracy^42^. Additionally, overall prompt length can counterintuitively decrease performance of LLMs for relatively simple tasks^43^. The additional context provided for minimum ornamentation depth may not have overcome the additional context size. This expert provided context, represented by the examples in the few-shot model, did improve accuracy for maximum and minimum wall thickness, and was at least statistically neutral in other cases.

Avoiding systematic bias is also important when using LLMs to extract data. The small local gemma3 model had an overall tendency to underestimate trait values (Figure 3). This improved with the larger Llama3.3 model. There has been a great deal of research detailing multiple kinds of bias in LLM models, and it is important to note that all LLM models will contain some degree of systematic bias^44^. Our results highlight the importance of quantifying both error rates and bias. When using LLMs to extract data from unstructured text, having a set of expert curated data is extremely helpful in generating benchmarks to assess these sources of error. Experts must supervise the use of LLMs in scientific endeavors, and benchmarking is a key way to utilize this expertise^45^.

One major advantage of LLMs over more traditional forms of natural language processing is the lack of reliance on feature engineering, the preprocessing of inputs into more useful data artifacts^46^. LLMs can ingest raw human generated text, and new models are even capable of ingesting images^47^. The ability to give LLMs raw text inputs with minimal preprocessing is a major advantage for analysis speed and therefore throughput. Conversely the unpredictability of these models mean they can struggle to meet the high accuracy of more traditional natural language processes techniques where precision greater than 95% is often possible^48^. Future work is needed to continue to improve the overall performance of LLM based data extraction, but the flexibility that this pipeline offers presents opportunities to generate very large datasets quickly. Researchers will continue to need to exercise prudence when balancing the needs of accuracy and efficiency, as well as rigor in validating the output of their chosen tools.

In conclusion, we demonstrate the potential of leveraging Large Language Models to accelerate the generation of fungal trait data from unstructured text resources. While accuracy varies by trait and model, and expert supervision remains essential, our pipeline offers a significant step towards overcoming the bottleneck of data scarcity in ecological research. The original creation of the TraitAM dataset involved numerous hours of manual labor by expert mycologists, a process that would be difficult to replicate for broader taxonomic groups. The ability to rapidly extract trait data from thousands of species descriptions opens doors to building comprehensive trait databases, enabling more robust ecological modeling and conservation efforts. This approach also provides a blueprint for applying LLMs to build trait databases across new traits and all domains of life.

Future work should focus on refining the prompt engineering and incorporating more sophisticated techniques to address challenges such as mathematical calculations and nuanced trait descriptions. Exploring multimodal approaches that integrate image recognition and other data sources could enhance the accuracy and richness of extracted information. Furthermore, the potential for combining LLMs with existing trait imputation methods warrants exploration^49–51^. LLMs could be used to generate preliminary trait estimates, with a small, expert-confirmed dataset used both to validate the LLM’s performance and to serve as a basis for imputation models. Automated routines could be used to flag potentially suspicious values, such as those exhibiting significant deviations from expected ranges or demonstrating unexpected combinations of traits, for subsequent expert confirmation. By leveraging the strengths of both approaches, we can move towards more comprehensive and accurate trait databases, ultimately driving ecological discovery and conservation efforts. As LLMs continue to evolve, we anticipate their increasing utility in transforming the way biological data is generated, managed, and utilized to address pressing ecological and conservation challenges.

